# Circular RNA *Pde4dip* regulates myogenesis by interacting with *Zfp143* mRNA: A novel regulatory axis

**DOI:** 10.1101/2025.09.17.676975

**Authors:** Suman Singh, Arundhati Das, Amaresh Chandra Panda

**Affiliations:** Institute of Life Sciences, Nalco Square, Bhubaneswar, Odisha, India

**Keywords:** RNA-RNA Interactions, circular RNA, mRNA regulation, Myogenesis

## Abstract

Many transcriptional and post-transcriptional regulators tightly regulate skeletal muscle myogenesis, including recently discovered circular RNAs (circRNAs). In this study, we used a crosslinking-based sequencing method called CliPP-Seq, wherein we performed AMT-mediated <u>C</u>ross-<u>Li</u>nking of RNA-RNA duplexes followed by <u>P</u>oly-A RNA <u>P</u>ulldown and sequencing to identify mRNA-interacting circRNAs in the mRNA samples of mouse C2C12 myoblasts and myotubes. BLAST analysis of the circRNAs with mRNAs identified their potential interacting partners. Interestingly, silencing of the circular RNA Pde4dip (*circPde4dip*) altered the target mRNA *Zfp143* (zinc finger protein 143) expression and suppressed the differentiation of C2C12 myoblasts into myotubes. In summary, we identified unexplored mRNA-interacting circRNAs and their possible role in muscle cell differentiation, specifically the *circPde4dip-Zfp143* mRNA interaction as one of the key regulators of myogenesis.

**Highlights:**

- Identified mRNA-associated circRNAs in differentiating C2C12 myoblasts
- The first report identifying the regulation of myogenesis by circRNA-mRNA interaction
- *circPde4dip* regulates C2C12 differentiation by interacting with *Zfp143* mRNA

## Introduction

The skeletal muscle forms the largest organ responsible for movement and metabolism [1]. Skeletal muscle is generated and maintained by the proliferation and differentiation of myogenic stem cells [2]. A range of transcription factors, including Pax3, Pax7, MyoD, Myf5, Myogenin, and Mrf4, are pivotal in regulating gene expression during the proliferation and differentiation of myoblasts to myotubes [3]. In addition to the well-studied transcription factors, recent research has suggested that non-coding RNAs, such as microRNAs (miRNAs), long non-coding RNAs (lncRNAs), and recently discovered circular RNAs (circRNAs), also play significant regulatory roles [4, 5]. CircRNAs have elicited considerable interest owing to their extraordinary stability, evolutionary conservation, tissue-specific expression, and altered expression in various developmental or disease states [6]. For instance, their expression levels are substantially altered during aging and muscle cell differentiation [7]. The involvement of circRNAs in myogenesis has only become apparent in recent years, and many studies have demonstrated that circRNAs regulate myogenesis mainly by interacting with miRNAs and RBPs. While most studies have reported the interaction of circRNAs with miRNAs or proteins, no clear picture of the direct circRNA-mRNA interaction has emerged so far [8].

Previously, we reported hundreds of circRNA-mRNA interactions in Beta-TC-6 cells, 2-days differentiated C2C12 cells, and HeLa cells [9]. Another report demonstrated that *circZNF609* directly binds to *CKAP5* mRNA and increases CKAP5 translation, regulating microtubule function in cancer cells and supporting cell-cycle progression [10]. Furthermore, to understand the role of mRNA-interacting circRNAs during myogenesis, we performed AMT-mediated crosslinking of transcripts expressed in C2C12 myoblast and 6-days differentiated myotube cells, followed by mRNA-hybrid complex pulldown and high-throughput sequencing called CLiPP-Seq [9]. We identified approximately 700 mRNA-interacting circRNAs with varying expression during myoblast differentiation. In this study, we investigated one such mRNA-interacting circRNA, *circPde4dip*, which most prominently interacts with *Zfp143* mRNA. We demonstrated that silencing *circPde4dip* led to increased expression of its interacting *Zfp143* mRNA and ZFP143 protein and reduced the differentiation of C2C12 myoblasts into myotubes. This study revealed a previously unknown regulatory role of circRNAs directly interacting with mRNA in myogenesis and emphasized the direct effect of circRNA on mRNA expression.

## Results

### CLiPP-Seq identified mRNA-interacting circRNAs in C2C12 myoblasts and myotubes

To understand the expression of mRNA-interacting circRNAs during myogenesis, C2C12 myoblasts, and 6-days differentiated C2C12 myotubes were used for AMT-treated CLiPP-sequencing. Analyzing the CLiPP-seq reads with the CIRCexplorer2 annotation, we identified 697 mRNA-interacting circRNAs in myoblasts and myotubes combined (**Supplementary Table S1, Figure 1A**). A majority of the circRNAs had a splice length of 500 nucleotides, while a few of the identified circRNAs had a splice length of > 1 kb (**Figure 1B**). circRNAs were derived from mouse chromosomes in an unbiased manner, represented along the mouse (mm10) cytoband (**Supplementary Figure 1A**). These data also showed that 70% of circRNAs were derived from exons, and only a part of the remaining fractions corresponded to circular intronic RNAs (ciRNAs) **(Figure 1C**). We also found that most circRNAs in CLiPP-seq were derived from genes with fewer than ten exons (**Supplementary Figure 1B**). Few host genes produced multiple circRNAs, but nearly all host genes were capable of producing one or two circRNAs (**Supplementary Figure 1C**). BLAST analysis identified that about 97% of circRNAs had complementarity with mRNAs in C2C12 CLiPP-seq datasets (**Supplementary Table S2, Figure 1D**).

**Figure 1.**
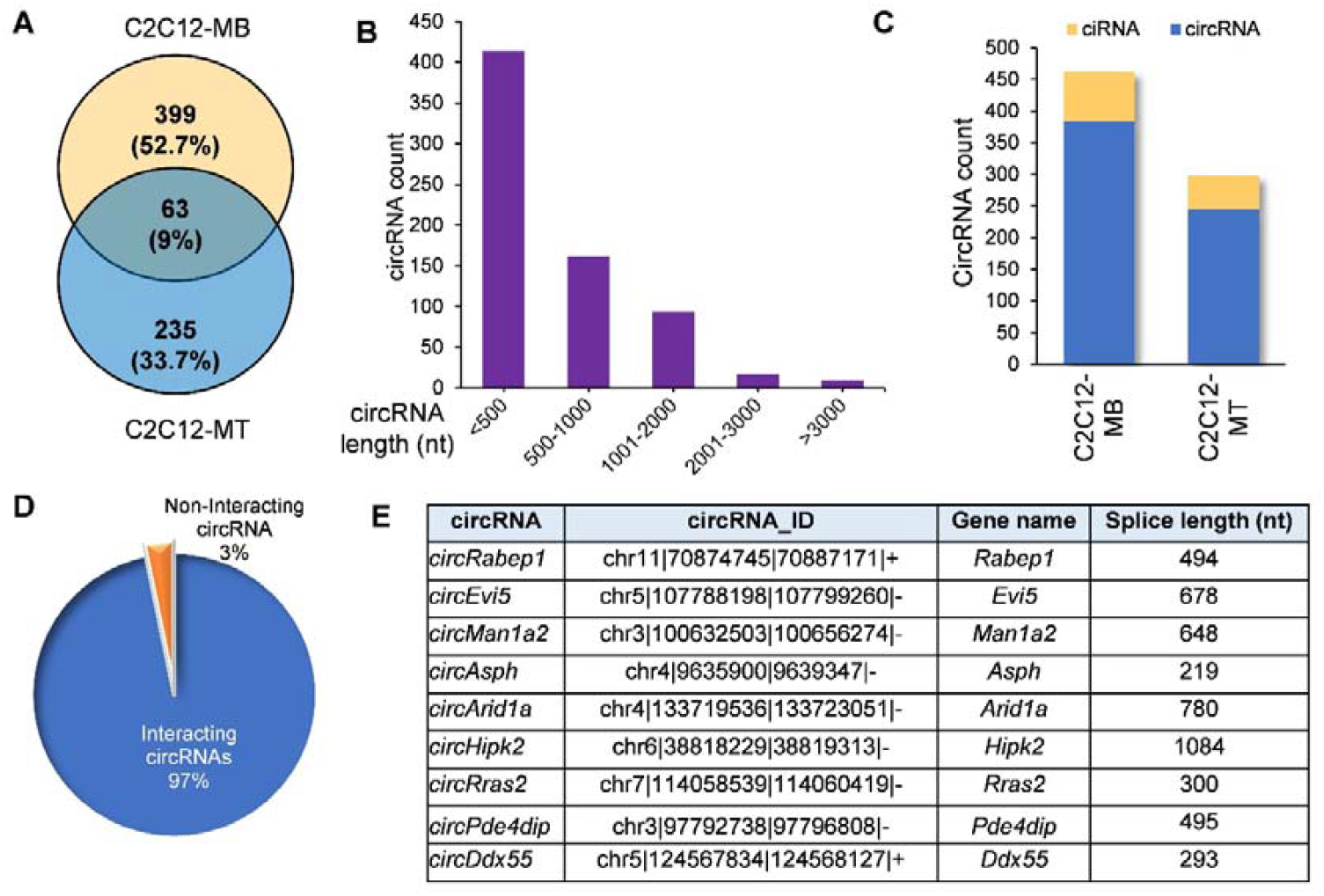
The mRNA-interacting circRNAs identified in C2C12-myoblast and myotube by CLiPP-Seq and their characteristics. **(A)** Venn diagram showing circRNAs identified in C2C12 myoblast and myotube-specific CLiPP-seq data sets. **(B)** Bar graph representing the splice lengths of total circRNAs identified in C2C12 myoblast and myotubes. **(C)** Bar graph showing exonic and intronic circRNAs identified in both datasets. **(D)** Pie chart showing the number of circRNAs interacting with mRNAs identified in BLAST analysis of CLiPP-seq samples. **(E)** Selected list of mRNA-interacting circRNAs in C2C12 myoblasts and myotubes.

### Characterization of differentiation-associated mRNA-interacting circRNAs

From the CLiPP-seq data, we shortlisted a few circRNAs based on a few criteria, including their splice length of <1.5 kb, exon count of <5, and abundance in C2C12 myoblasts or myotube cells **(Figure 1E)**. We used divergent primer pairs across the backsplice junction to validate the expression of mRNA-interacting circRNAs in C2C12 cells by RT/ no-RT followed by PCR (**Figure 2A**). As shown in **Figure 2B**, purification of the amplified PCR products and subjecting them to Sanger sequencing proved that only the backsplice junction sequences of target circRNAs were specifically amplified. We further analyzed the target linear or circRNAs in total RNA from C2C12 cells treated with RNase R, as it is well known that circRNAs are resistant to exonuclease RNase R owing to the absence of free ends. Linear *Actb* and *Gapdh* mRNAs were significantly degraded by RNase R digestion, whereas the selected mRNA-interacting circRNAs were immune to degradation, confirming their identity as circular transcripts **(Figure 2C)**.

**Figure 2.**
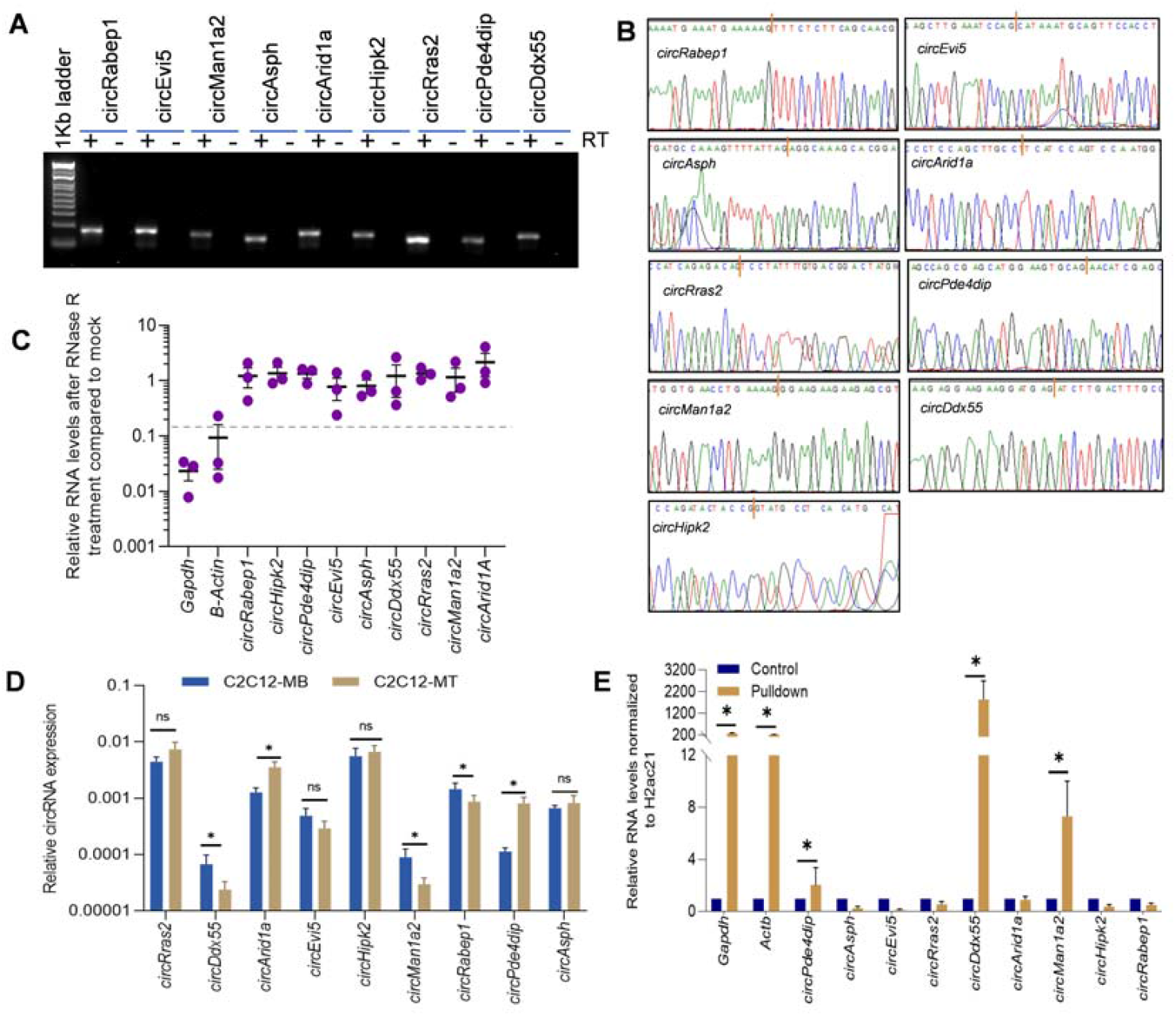
Validation of circRNAs and their enrichment in mRNA pulldown. **(A)** Visualization of RT/no-RT PCR amplicons in SYBR green stained 2% agarose gel. **(B)** The chromatogram shows the circRNA sequence, and the red bar specifies the junction sequences. **(C)** Interleaved scatter plot showing qRT-PCR analysis after RnaseR treatment. (**D)** RT-qPCR analysis showing the expression pattern of circRNAs in C2C12-myoblast compared to C2C12-myotube stage. **(E)** RT-qPCR analysis showing enrichment of *circPde4dip, circDdx55*, and *circMan1a2* along with control *Gapdh* and *Actb* mRNAs upon oligo d(T) mediated mRNA pulldown.

### mRNA-interacting circRNAs are differentially expressed during myogenesis

Proliferating C2C12 myoblast and 4-days differentiated C2C12 myotubes were used to analyze the differential expression of selected circRNAs. Microscopic images and upregulation of myogenesis markers confirmed the differentiation of C2C12 myoblasts into myotubes (**Supplementary Figure 2A, B)**. Interestingly, some of the selected mRNA-interacting circRNAs, including *circDdx55, circArid1a, circMan1a2, circPde4dip*, and *circRabep1* showed significant differential expression patterns in C2C12 myotubes compared to myoblasts **(Figure 2D)**. Furthermore, we performed a poly(A) RNA pulldown assay using oligo-dT beads with total RNA from 2-days differentiated C2C12 myotubes, followed by enrichment analysis to check the interaction of circRNAs with poly(A) mRNAs/mRNAs. Notably, several circRNAs, including *circPde4dip, circDdx55*, and *circMan1a2*, along with *Gapdh* and *Actb* mRNA, were significantly enriched compared to the input samples, confirming the specific pulldown of poly(A) RNAs and their interacting circRNAs (**Figure 2E)**. As *circPde4dip* expression showed the highest upregulation in myotubes compared to myoblasts and significant enrichment in poly(A) RNA pulldown (**Figure 2D, E**), we selected *circPde4dip* to further extrapolate its role during C2C12 differentiation.

### *circPde4dip* is stably expressed and interacts with multiple mRNA targets

Back splicing of *Pde4dip* pre-mRNA (ENSMUSG00000038170) produces a ∼495-nucleotide exonic circular *Pde4dip* RNA (**Figure 3A, Supplementary Figure 3A**). Analysis of the stability of *circPde4dip* in 2-days differentiated C2C12 cells with actinomycin D treatment indicated that *circPde4dip* remained stable until 8 h post-treatment, while *Myc* mRNA progressively degraded within an hour **(Figure 3B)**. We analyzed the BLAST data to identify several mRNAs that interact with *circPde4dip* in C2C12 cells, as depicted in Cytoscape (**Figure 3C, Supplementary Figure 3B)**. Since *circPde4dip* was enriched in C2C12 myotubes in the CLiPP-seq data, we checked the expression pattern of its interacting mRNAs as well in C2C12 myoblasts and myotubes. RT-qPCR analysis confirmed reduced expression of *circPde4dip-*interacting zinc finger protein 143 (*Zfp143)* and *Tsc22d* mRNA in myotubes compared to myoblasts (**Figure 3D)**. Acknowledging the potential for false-positive results in poly(A) pulldown from total RNA due to mere sequence complementarity, we performed a native pulldown assay on 2-days differentiated C2C12 cell lysates. Notably, the enrichment of *circPde4dip* along with *Gapdh* and *Actb* mRNA from poly(A) pulldown in the cell lysates further indicated a likely interaction with target mRNAs (**Figure 3E)**. Further, using the inverse approach, *circPde4dip* was pulldown in 3-days differentiated C2C12 cell lysates using biotin-tagged antisense oligos (ASO) targeting the backsplice junction sequence, followed by RT-qPCR analysis, which showed significant enrichment of *circPde4dip* along with the target *Zfp143* mRNA, indicating direct base-pairing with *circPde4dip* (**Figure 3F**).

**Figure 3.**
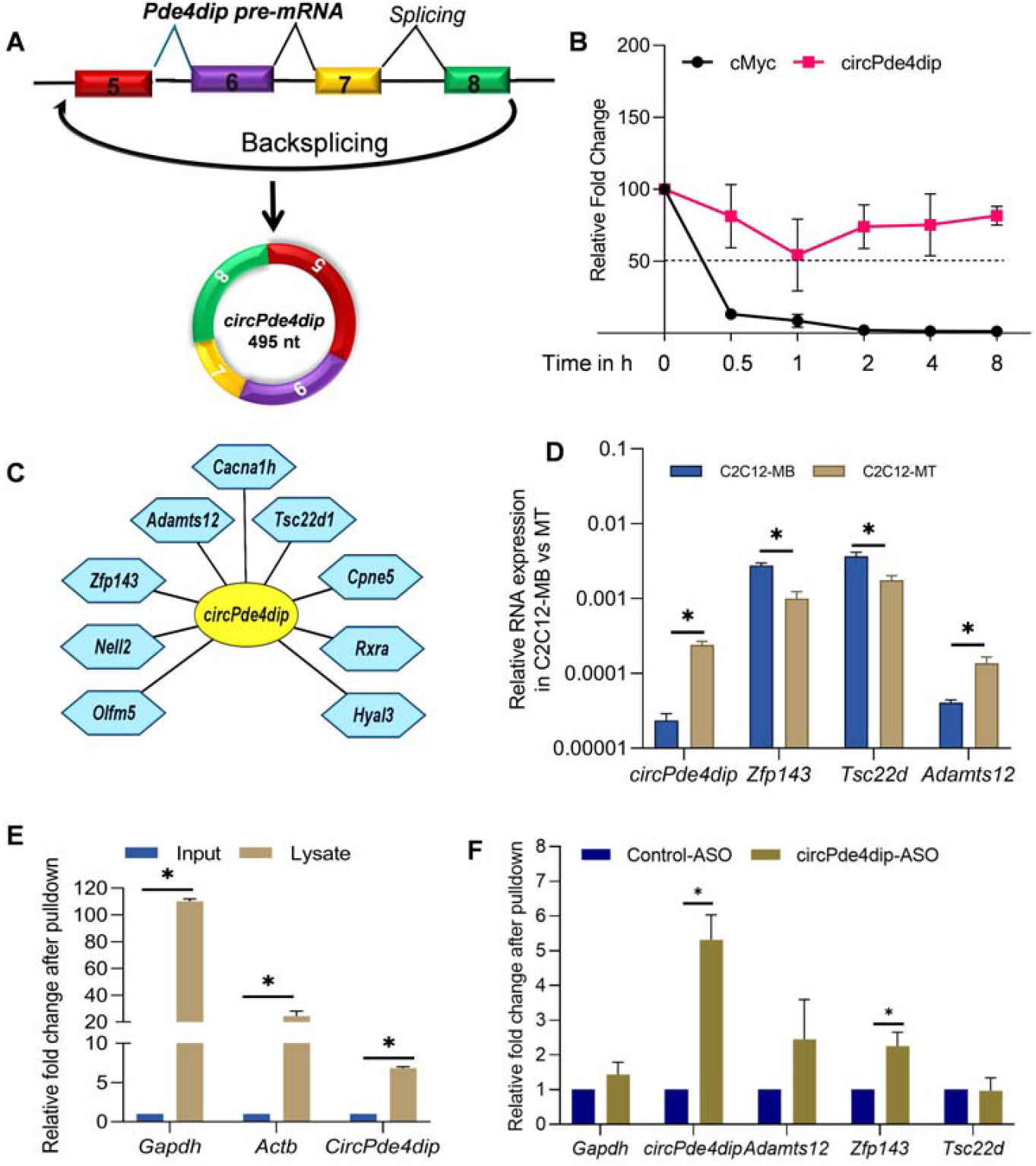
Validation of *circPde4dip*-interacting mRNA expression in C2C12. (**A**) Schematic representation of circPde4dip biogenesis via backsplicing **(B)** RT-qPCR analysis showing relative fold change in *circPde4dip* expression compared to linear *Myc* mRNA after Actinomycin D treatment. (**C)** Cytoscape showing *circPde4dip* specific mRNA targets based on sequence complementarity. **(D)** RT-qPCR analysis showing the differential expression of *circPde4dip-*interacting mRNAs in C2C12 myoblast compared to myotube stage. **(E)** RT-qPCR analysis showing the enrichment of *circPde4dip* in poly(A) pulldown samples compared to input cell lysates of 2-days differentiated C2C12 cells. **(F)** RT-qPCR result showing enrichment of *Zfp143* mRNA along with *circPde4dip* upon ASO pulldown.

### *circPde4dip* regulates *Zfp143* expression and promotes myogenesis in C2C12 cells

Since *circPde4dip* expression was upregulated at the myotube stage, we hypothesized that *circpde4dip* could regulate C2C12 muscle cell differentiation. We silenced *circPde4dip* using a backsplice junction-specific GapmeR and allowed it to differentiate for three days. The phase-contrast images showed reduced myotube formation upon *circPde4dip* silencing compared to control C2C12 cells (**Figure 4A**). Knockdown of *circPde4dip* led to a significant increase in *Zfp143* mRNA expression, whereas *Tsc22d* and *Adamts12* mRNA expression was not affected, indicating that the regulation of myogenesis by *circPde4dip* could be mediated through *Zfp143* (**Figure 4B)**. Further, we observed that ZFP143 protein expression was upregulated upon *circPde4dip* silencing, indicating direct regulation of ZFP143 by *circPde4dip* (**Figure 4C)**. Furthermore, to rule out miRNA-mediated indirect regulation of *Zfp143* mRNA, we performed miRNA analysis of *circPde4dip* targets and, interestingly, found that *Zfp143* mRNA was not targeted by any of the *circPde4dip*-associated miRNAs (**Supplementary Figure 3C, D**). Using bimolecular structure predictions and pulldown assays, we proved that circRNAs might directly hybridize with mRNAs to form RNA duplexes (**Supplementary Figure 4**). Our results demonstrated that increased expression of ZFP143 upon *circPde4dip* silencing could promote the cells to maintain their stemness and hence prevent myotube formation **(Figure 4D)**.

**Figure 4.**
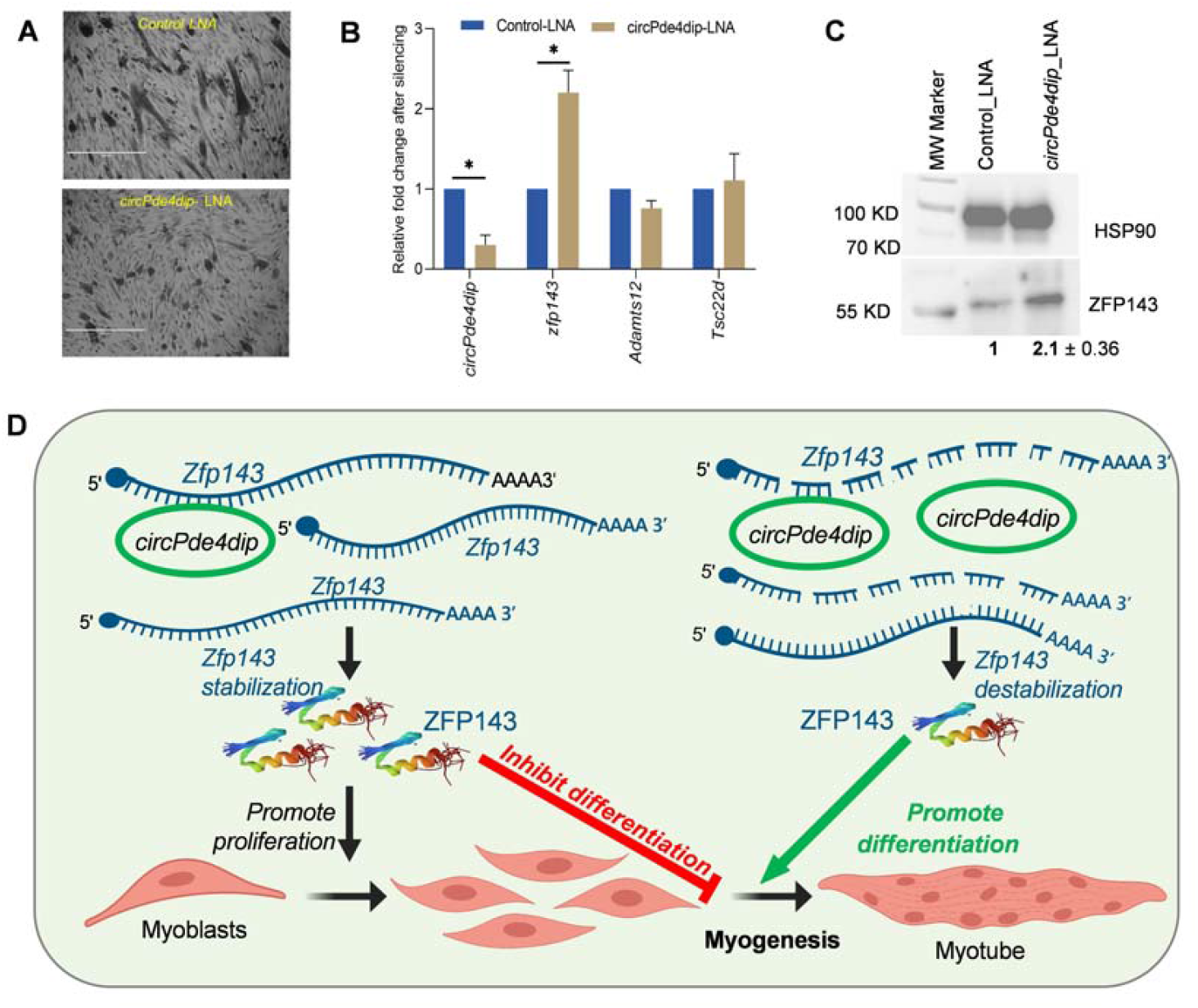
Regulation of C2C12 differentiation by *circPde4dip*. (**A**) The phase contrast image showing C2C12 myotubes, 72hr post treatment with control LNA and *circPde4dip* LNA. **(B)** RT-qPCR analysis showing relative fold change in *circPde4dip* expression along with target mRNA expression after silencing. **(C)** Western-blot showing ZFP143 expression compared to control HSP90 expression after *circPde4dip* silencing. **(D)** Possible mechanism of C2C12 differentiation mediated by *circPde4dip* interaction with *Zfp143*. In the myoblast stage the inhibition of C2C12 cells differentiation is driven by the presence of *Zfp143* mRNA which encodes ZFP143 transcription factor responsible for maintaining the cells at myoblast stage by promoting proliferation. In the myotube stage, *circPde4dip* destabilizes its interacting partner *Zfp143* mRNA leading to reduced expression of ZFP143 and promoting myotube formation.

## Discussion

CircRNAs have been proven to be key regulators of the pre-mRNA splicing machinery, transcription, translation, and protein functions by interacting with target miRNAs or RBPs. Given the diverse roles of circRNAs, there has been a significant increase in reports suggesting role of circRNAs in skeletal muscle differentiation [5]. The ability of circRNAs to regulate myogenesis has been mostly limited to their miRNA sponging activity, with a few reports on circRNA-RBP interactions in myogenesis. The interplay between circRNAs and mRNAs remains largely unexplored. In this study, we investigated the circRNA-mRNA crosstalk in regulating myogenesis.

Recent shift in research highlights the role of direct interaction of circRNAs with mRNAs, evidenced by a novel finding showing circZNF609 binds to CKAP5 mRNA and thus plays an important role in regulating microtubule faltering and tumorigenesis [11]. Previously, we reported the direct interaction of circRNAs with mRNAs in HeLa, βTC6, and C2C12 cells [9]. Another study report on circRNA-mediated mRNA decay further strengthens our hypothesis wherein the circRNA-mRNA interaction resulted in bringing the exon-junction complex (EJC) into proximity with the 3’UTR, leading to EJC-dependent nonsense-mediated mRNA decay (NMD) [12]. From our BLAST analysis, we understand that most circRNA interactions occur in the UTR region of mRNAs, highlighting their potential regulatory significance [9]. This study is our further attempt to understand the differential roles of mRNA-interacting circRNAs in muscle cell differentiation. From the CLiPP-Seq data, a large number of mRNA-interacting circRNAs were identified in C2C12 myoblasts and myotubes (**Figure 1)**. Furthermore, from the BLAST analysis, we found that 97% of the circRNAs interacted with mRNA. From the pulldown assays, we determined that *circPde4dip* was significantly enriched with linear mRNAs, and from the differential expression analysis in C2C12 by RT-qPCR, we found *circPde4dip* to be an interesting target, as it was significantly upregulated during C2C12 cell differentiation (**Figure 2**). Although many mRNAs were predicted to be complementary to *circPde4dip*, only *Zfp143* mRNA was significantly enriched upon *circPde4dip* pulldown (**Figure 3**).

Aligning with the reported findings, we found that *Zfp143* mRNA was downregulated in the differentiated C2C12 cells, which showed an inverse expression pattern in the C2C12 myotubes compared to *circPde4dip* (**Figure 3**). Furthermore, an interesting inverse correlation between ZFP143 and *circPde4dip* expression was observed during myogenesis and circRNA silencing (**Figure 4**). These findings suggest *circPde4dip* interacts with *Zfp143* mRNA directly, which might lead to destabilization of mRNA during the differentiation process. Notably, circRNA of interest, *circPde4dip*, was found to specifically bind to the 5’ UTR of *Zfp143* mRNA. The binding of circRNA to the 5’-UTR could obstruct its crucial role in ribosome recruitment, which could lead to stalled translation and subsequent mRNA decay. Alternatively, circRNA binding might induce secondary structure alterations in the 5’UTR that could expose mRNA to stress-induced degradation. Additionally, the competitive binding of circRNAs in the 5’UTR could potentially disrupt the association of the cap-binding complex (CBC) and reduce mRNA stability. *Zfp143* mRNA has been previously reported to maintain the pluripotency in mouse embryonic stem cells [13] and is a major regulator in rat vascular smooth muscle cells [14]. As ZFP143 promotes stemness and cell proliferation, *circPde4dip* silencing leads to reduced myotube formation due to the upregulation of ZFP143. Together, we hypothesized that *circPde4dip* inhibits the expression of ZFP143 by interacting directly with the mRNA. In the differentiation process, upregulated *circPde4dip* interacted with *Zfp143* mRNA, leading to its destabilization and suppression of ZFP143, ultimately favoring myotube formation **(Figure 4)**.

Despite our work confirming the direct regulation of gene expression by circRNA-mRNA interactions in skeletal muscle, there are few considerable limitations. First, the number and types of circRNAs could be different based on the computational pipeline and sequencing depth used for circRNA identification. Second, the positions of circRNA–mRNA interactions were computationally identified. Third, considering that circRNAs and mRNAs interact with RBPs as well as miRNAs, whether the interacting sequences are free to interact with each other and which interacting-RBP can indirectly regulate target mRNA requires further experimental validation. Fourth, the *in vivo* secondary and tertiary structures of interacting RNAs in endogenous cellular conditions may need to be considered when dissecting the functional significance of circRNA-mRNA hybrids. While these circRNA-mRNA mechanisms present intriguing possibilities, an in-depth study of this regulatory mechanism can help us broaden our understanding of the mRNA-interacting circRNAs in myogenesis and other cellular events. Our study not only discovers novel links between hundreds of circRNAs and mRNAs but also highlights the need for follow-up work that can provide a considerable scope for the development of new RNA-based treatments.

### Experimental procedures

#### Cell culture and differentiation

Mouse C2C12 myoblast cells were cultivated in a growth medium consisting of Dulbecco’s Modified Eagle’s Medium (DMEM) with Fetal Bovine Serum (FBS) of 15% concentration and penicillin-streptomycin (1%). The growth medium of sub-confluent myoblast cells was replaced with a differentiation medium containing DMEM supplemented with 2% horse serum and 1% penicillin-streptomycin [15]. The cells were supplemented with differentiation media for up to six days depending on the requirement, and differentiation was visualized under a phase-contrast microscope to monitor the formation of multinucleated elongated myotubes.

#### AMT-mediated proximity ligation and RNA sequencing

The 4′-aminomethyltrioxsalen hydrochloride (AMT), a compound known for its ability to induce RNA-RNA crosslinking, was diluted in DMEM and 2X phosphate buffer saline to obtain a working concentration of 20 μg/ml, which was used as the irradiation media. C2C12 myoblast cells and 6-days differentiated C2C12 myotubes were placed in irradiation media and then incubated at 37°C for 30 min [16]. UV crosslinking was then conducted at 365 nm to facilitate RNA-RNA crosslinking [17].

Total RNA was isolated from AMT-treated cells, followed by quality check using NanoDrop 2000, Multiskan Sky, or Qubit 4. Subsequently, the mRNA fraction was purified from 1μg of the total RNA using the NEBNext® poly(A) mRNA Magnetic Isolation Module (NEB), followed by concentration and quality checks. The mRNAs and its interacting RNA complexes pulled down from the C2C12 cells were used for Crosslinking-Immunoprecipitation-Sequencing (CLiPP-Seq). The cDNA libraries were prepared using the NEB NeXT Ultra II Library Prep Kit (NEB) with minor modifications to incorporate additional steps of de-crosslinking at 254 nm after RNA fragmentation [16]. The cDNA libraries were then analyzed on TapeStation, followed by 150 bp paired-end sequencing on the Illumina HiseqX platform.

#### RNA sequencing analysis and BLAST

The adapters from the RNA-seq reads were removed using the Cutadapt tool, and quality was checked by Fastqc, followed by alignment with the mouse reference genome assembly release of mm10. For circRNA identification, reads were aligned using the STAR aligner using the ChimSegmentMin-10 parameter of the CIRCexplorer2 pipeline [18]. The mature sequences of the circRNAs were used to perform BLAST against the mouse transcript (NCBI BLAST), as well as command line BLAST, and only Plus/Minus interactions were filtered to identify the circRNA-mRNA hybrids.

#### RNA isolation, RT-PCR, Sanger sequencing, and Quantitative (q)PCR

Total RNA was extracted from 2-days differentiated C2C12 cells using either the TRIzol method or the MagSure All RNA isolation kit (RNA Biotech). cDNA synthesis with or without RT enzyme (+/-RT) was performed from total RNA using High Capacity or Maxima Reverse Transcription Kit (Thermo). The RT/no-RT cDNA samples were then amplified with divergent primers, using 2X DreamTaq PCR master mix, followed by visualization of the PCR amplicons on a 2% SYBR gold-stained agarose gel. For backsplice junction sequence validation, RT-PCR products were purified and subjected to Sanger sequencing using one of divergent primers [19]. For quantitative PCR, 2X PowerUP SYBR Green Master Mix (Thermo) was used, and target enrichment was calculated using the delta-CT method, considering *18s* rRNA or *Gapdh* mRNA as internal controls [20].

#### RNase R treatment and stability assay of circRNA & mRNA

For RNase R treatment assay, total RNA was isolated from 2-days differentiated C2C12 cells, 5 μg of which was treated with or without 0.5 μL of RNase R (Lucigen) at 37°C for 60 min, followed by RNA isolation and cDNA synthesis. RT-qPCR analysis was performed on mock and RNase R-treated samples using specific primers for linear and circRNAs (**Supplementary Table S3**) [21]. To check the stability of circRNAs over mRNAs, 2-days differentiated C2C12 cells were treated with actinomycin D (10 μg/mL) for 0h, 0.5, 1, 2, 4, and 8 h, followed by total RNA isolation, cDNA synthesis, and RT-qPCR analysis of *circPde4dip* and *c-Myc* expression levels at different time points.

#### Poly(A) RNA and circRNA pulldown assays

For mRNA pulldown, 10μg of the total RNA isolated from 2-days differentiated C2C12 cells was subjected to poly(A) pulldown using NEBNext® Poly(A) mRNA Magnetic Isolation Module (NEB) following the manufacturer’s instructions. For native mRNA pulldown assay, 2-days differentiated C2C12 cells were lysed with polysome extraction buffer (PEB), followed by incubation with NEBNext Oligo d(T)25 Beads in 1XTENT buffer for 90 min at 4°C in a rotator. The beads were then washed with the wash buffer, followed by elution of mRNA in 12 μl of elution buffer. One μg of total RNA was used as an input control and total 12 μl of eluted mRNA was used for cDNA synthesis and RT-qPCR analysis of mRNA and circRNAs in pulldown samples compared to input samples [9]. circRNA pulldown was performed using biotin-labelled antisense oligo (ASO), as described previously [22]. Briefly, 3-days differentiated C2C12 cells were lysed with polysome extraction buffer (PEB). Subsequently, the supernatant was collected and equally distributed for pulldown with control-ASO or *circPde4dip*-ASO. An equal volume of (2X) TENT buffer was then added to the supernatant and subjected to hybridization using 1 μl of 100 μM control-ASO and *circPde4dip*-ASO, followed by ASO-complex pulldown using streptavidin beads. Both control and circRNA pull-down samples were used for cDNA synthesis and RT-qPCR analysis.

#### *circPde4dip* silencing and western blotting

For circRNA silencing, antisense GapmeR oligos targeting the unique backsplice junction sequence of circRNAs were synthesized. Actively growing C2C12 cells were subjected to GapmeR LNA transfection with 100 nM twice at 24 h intervals using Lipofectamine RNAiMAX (Invitrogen) and allowed to differentiate for 72 h. Total RNA was then isolated, followed by cDNA synthesis and RT-qPCR analysis to determine the expression of circRNA and target mRNA. Using the same experimental design, 72-hour post-transfection total cellular protein was isolated using RIPA buffer (Himedia), followed by western blot analysis of ZFP143 and HSP90 expression.

### Statistical analyses and visualization

In the present study, Student’s t-test was employed to calculate the statistical significance of the observed findings, with a threshold of *P < 0.05, as statistically significant. All experimental results and figures were obtained from at least three independent biological replicates, ensuring the reliability and robustness of the results. For data visualization and analysis, various software tools were utilized, including Microsoft Excel, GraphPad Prism, Cytoscape, and R studio.

## DECLARATIONS

### Data availability

All the data generated in this study are included in the main text or supplementary data. The RNA-seq data generated in this study were deposited in the Indian Biological Data Centre (IBDC) with INSDC Project Accession number PRJEB90311/ERP173326. The Supplementary Tables can be found in the Figshare data repository (https://figshare.com/s/92d62b495736fe07100f)

### Author contributions

Suman Singh: Conceptualization, Methodology, Investigation, Formal analysis, Validation, Visualization, Writing—original draft, review, and editing. Aundhati Das: Methodology, Investigation, Writing-Review and Editing. Amaresh C. Panda: Conceptualization, funding acquisition, Supervision, Writing, review, and editing

## Acknowledgements

We thank Dr. Sharmishtha Shyamal for her help in uploading the RNA-seq data to IBDC and Mr. Gaurahari Sahoo for technical support in some experiments. We thank our colleagues for their helpful discussions and manuscript proofreading.

## Consent for publication

All authors consent this manuscript for publication.

## References

1. Stump, C.S., et al., The metabolic syndrome: role of skeletal muscle metabolism. Ann Med, 2006. 38(6): p. 389–402.

2. Frontera, W.R. and J. Ochala, Skeletal Muscle: A Brief Review of Structure and Function. Calcified Tissue International, 2015. 96(3): p. 183–195.

3. Buckingham, M., Gene regulatory networks and cell lineages that underlie the formation of skeletal muscle. Proceedings of the National Academy of Sciences, 2017. 114(23): p. 5830–5837.

4. Butchart, L.C., et al., The long and short of non-coding RNAs during post-natal growth and differentiation of skeletal muscles: Focus on lncRNA and miRNAs. Differentiation, 2016. 92(5): p. 237–248.

5. Das, A., et al., Circular RNAs in myogenesis. Biochim Biophys Acta Gene Regul Mech, 2020. 1863(4): p. 194372.

6. Voellenkle, C., et al. Dysregulation of Circular RNAs in Myotonic Dystrophy Type 1. International Journal of Molecular Sciences, 2019. 20, DOI: 10.3390/ijms20081938.

7. Lin, Z., et al., Functions and mechanisms of circular RNAs in regulating stem cell differentiation. RNA Biology, 2021. 18(12): p. 2136–2149.

8. Zhang, P., et al., Assessment of myoblast circular RNA dynamics and its correlation with miRNA during myogenic differentiation. Int J Biochem Cell Biol, 2018. 99: p. 211–218.

9. Singh, S., et al., Global identification of mRNA-interacting circular RNAs by CLiPPR-Seq. Nucleic Acids Res, 2024. 52(6): p. e29.

10. Rossi, F., et al., Circular RNA ZNF609/CKAP5 mRNA interaction regulates microtubule dynamics and tumorigenicity. Mol Cell, 2022. 82(1): p. 75–89.e9.

11. Rossi, F., et al., Circular RNA ZNF609/<em>CKAP5</em> mRNA interaction regulates microtubule dynamics and tumorigenicity. Molecular Cell, 2022. 82(1): p. 75–89.e9.

12. Boo, S.H., et al., Circular RNAs trigger nonsense-mediated mRNA decay. Mol Cell, 2024. 84(24): p. 4862–4877.e7.

13. Ye, J. and R. Blelloch, Regulation of pluripotency by RNA binding proteins. Cell Stem Cell, 2014. 15(3): p. 271–280.

14. Hernández-Negrete, I., et al., Adhesion-dependent Skp2 transcription requires selenocysteine tRNA gene transcription-activating factor (STAF). Biochem J, 2011. 436(1): p. 133–43.

15. Clemente, C.F., et al., Differentiation of C2C12 myoblasts is critically regulated by FAK signaling. Am J Physiol Regul Integr Comp Physiol, 2005. 289(3): p. R862–70.

16. Lu, Z., et al., RNA Duplex Map in Living Cells Reveals Higher-Order Transcriptome Structure. Cell, 2016. 165(5): p. 1267–1279.

17. Engreitz, Jesse M., et al., RNA-RNA Interactions Enable Specific Targeting of Noncoding RNAs to Nascent Pre-mRNAs and Chromatin Sites. Cell, 2014. 159(1): p. 188–199.

18. Dong, R., et al., Genome-Wide Annotation of circRNAs and Their Alternative Back-Splicing/Splicing with CIRCexplorer Pipeline. Methods Mol Biol, 2019. 1870: p. 137–149.

19. Singh, S., A. Das, and A.C. Panda, Sanger Sequencing to Determine the Full-Length Sequence of Circular RNAs, in Circular RNAs, C. Dieterich and M.-L. Baudet, Editors. 2024, Springer US: New York, NY. p. 93–105.

20. Panda, A.C. and M. Gorospe, Detection and Analysis of Circular RNAs by RT-PCR. Bio Protoc, 2018. 8(6).

21. Das, A., D. Das, and A.C. Panda, Validation of Circular RNAs by PCR. Methods Mol Biol, 2022. 2392: p. 103–114.

22. Das, D., A. Das, and A.C. Panda, Antisense Oligo Pulldown of Circular RNA for Downstream Analysis. Bio Protoc, 2021. 11(14): p. e4088.

