## Supplementary Figure for "Circular RNA *Pde4dip* regulates myogenesis by interacting with *Zfp143* mRNA: A novel regulatory axis"

### SUPPLEMENTARY TABLES AND FIGURES

**Supplementary Table S1:** CircRNA annotation of C2C12-seq

**Supplementary Table S2:** CircRNA-mRNA BLAST analysis in C2C12 cells

**Supplementary Table S3:** Oligonucleotides used in this study

**Singh et al, Supplementary Figure 1**

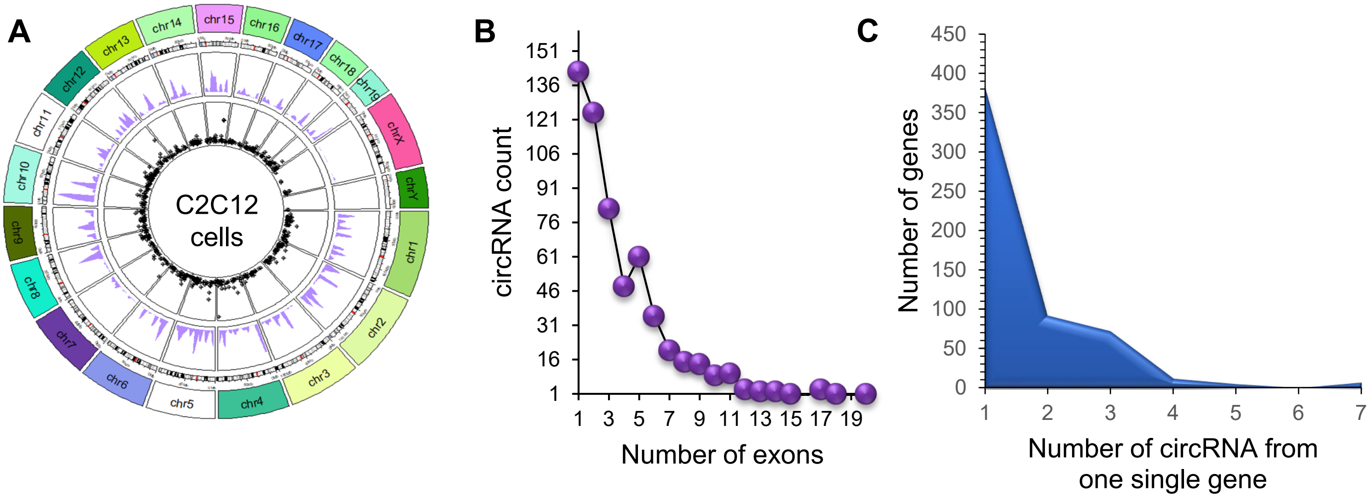


**Supplementary Figure 1. The mRNA-interacting circRNAs identified in C2C12 CLiPP-Seq data. (A)** Circos plotting the distribution of number and lengths of circRNAs of both datasets along the mouse ctyoband. **(B)** Scatter plot showing the exon retention of total circRNAs identified in MB and MT. **(C)** Area plot showing the number of circRNAs generated from each gene.

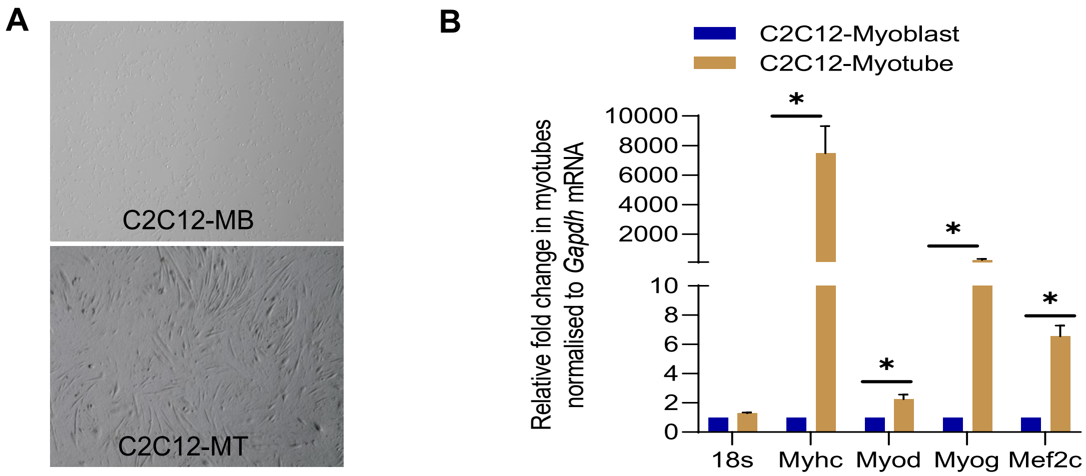


**Supplementary Figure 2. Validation of circRNA enrichment upon mRNA pulldown.** (**A)** The phase contrast field image showing C2C12 myoblast and day-4 differentiated myotubes. **(B)** RT-qPCR analysis showing the expression pattern of myogenic regulatory factors in C2C12 myotubes compared to proliferating myoblasts normalized to *Gapdh* mRNA. The error bars represent means ± SEM from 3 independent experiments, and * indicates the statistical significance with p-value < 0.05.

**A** circPde4dip splice sequence (495 nt)
AACATCGAGCTGAAGGTTGAAGTGGAGAGCCTGAAACGAGAACTCCAGGACAGGAAAC
AGCATCTAGATAAAACATGGGCCGATGCAGAGGATCTCAACAGCCAGAATGAGGCAGA
GCTCCGGCGCCAGGTTGAAGAACGGCAGCAGGAGACAGAACACGTTTATGAGCTCCTA
GGGAACAAGATCCAGCTGCTGCAGGAGGAACCCAGGCTAGCAAAGAATGAAGCCACA
GAGATGGAGACTCTGGTGGAGGCAGAGAAGAGGTGCAATCTGGAGCTCTCAGAGAGG
TGGACGAATGCTGCCAAGAACAGGGAAGATGCAGCAGGAGACCAGGAGAAGCCTGAC
CAATATTCTGAGGCACTGGCTCAGAGGGACAGGAGAATTGAAGAGCTGAGGCAGAGCT
TGGCTGCTCAGGAGGGGCTTGTGGAACAGCTGTCTCAAGAGAAACGACAACTGTTACA
TCTGCTGGAGGAGCCAGCGAGCATGGAAGTGCAG

**B**

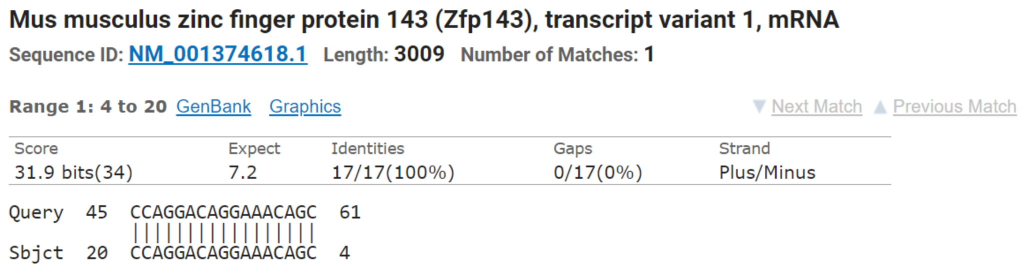


**C**

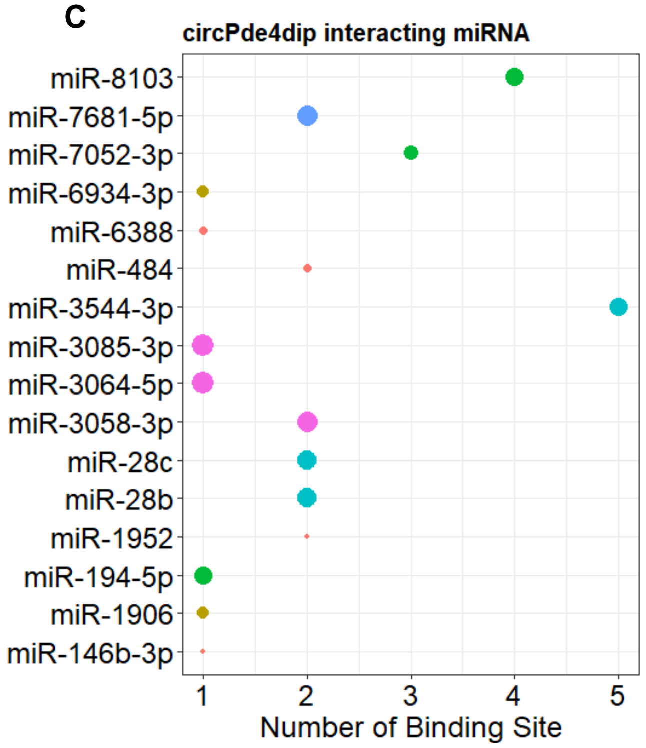


**D**

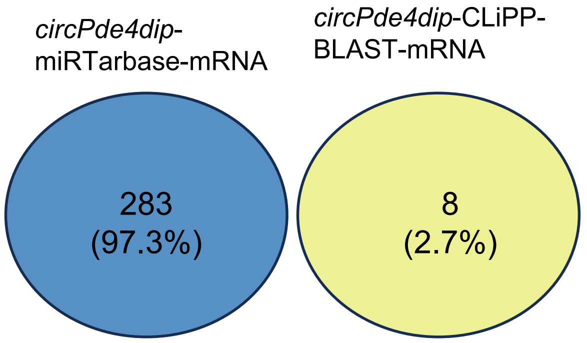


**Supplementary Figure S3.** *(***A)** The miRNA regulatory axis of *circPde4dip* **(B)** Screenshot of NCBI BLAST of *circPde4dip and Zfp143* mRNA. **(C)** Dot plot showing the number of binding sites and target scores of miRNAs predicted by miRDB to interact with *circPde4dip*. **(D)** Venn diagram showing the miRTarBase-predicted mRNA targets of miRNAs associated with *circPde4dip* and the mRNA targets of *circPde4dip* identified by BLAST in C2C12 CLiPP-seq datasets.

**Singh et al, Supplementary Figure 4**

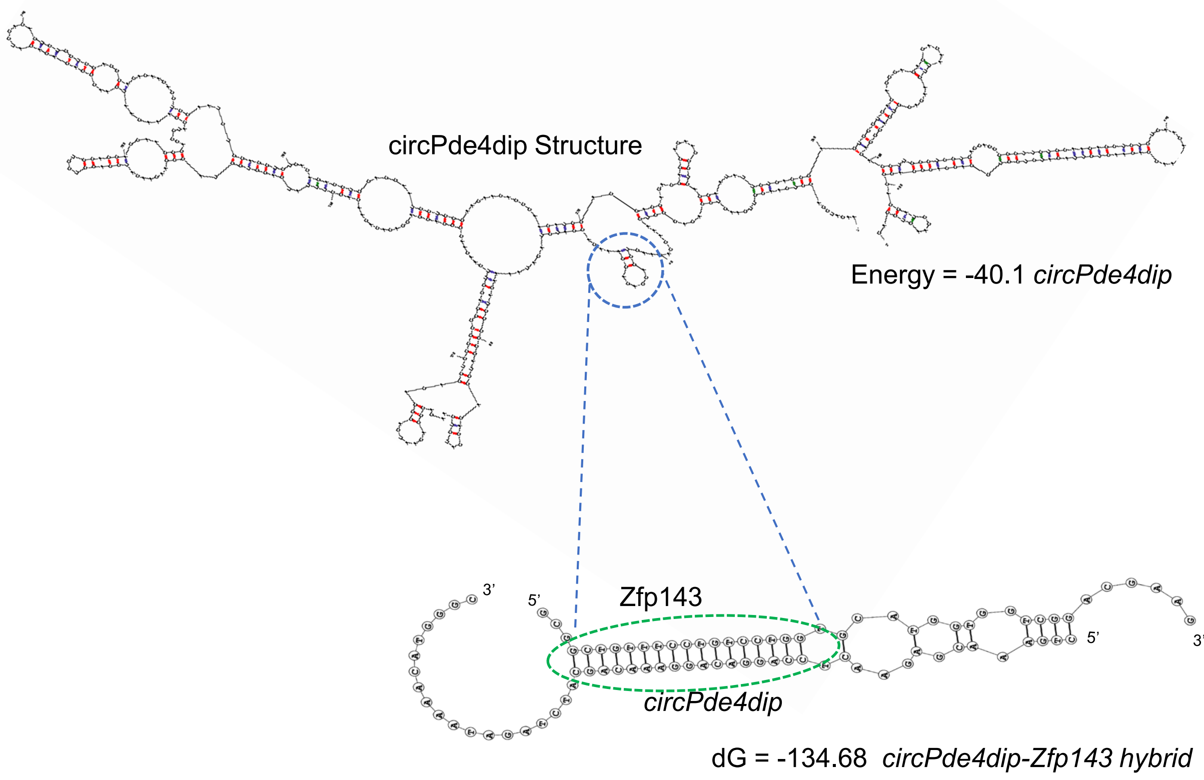


**Supplementary Figure S4:** Secondary structure of *circPde4dip* predicted by RNAstructure web server. The blue dotted circles indicate the region of interaction in *circPde4dip.* The green dotted circles indicate the bimolecular secondary structure and region of interaction of *circPde4dip* with the interacting sequences of target *Zfp143* mRNA predicted by the DuplexFold algorithm of RNAstructure web server.
